# Rational design of Cas9 ribonucleoprotein with a “gRNA-shRNA” for multidimensional genome manipulation and enhanced homology-directed repair

**DOI:** 10.1101/2022.03.17.484717

**Authors:** Jie Qiao, Wenli Sun, Wenhao Yin, Lixin Ma, Yi Liu

## Abstract

Gene perturbation approaches have evolved as powerful tools for understanding the function of genes and curing inherited diseases. Here, we develop a method that combines the merits of RNAi and CRISPR technology by rational design of Cas9 ribonucleoprotein (RNP) with a “gRNA-shRNA” component. The RNP, termed Cas9-RNAi, has a gRNA containing a 3’ extension that can be processed to a functional siRNA via dorsha/dicer enzyme mediated cleavage within cells. We prepared the Cas9-RNAi RNPs by streamline co-expression of Cas9 enzymes and the “gRNA-shRNA” ribonucleotides in *Escherichia coli* strain HT115(DE)3. Transferring the Cas9-RNAi RNPs into mammalian cells achieves multidimensional genome manipulation, e.g., simultaneously knock out and knock down the target genes. Moreover, by introduction of a shRNA against the gene of human DNA ligase 4 (*LIG4*), significantly improved homology-directed repair was attained. Together, we develop a simple-to-use CRISPR RNP tool that has great potentials in precise genome editing, gene function analysis and gene therapy.

## INTRODUCTION

The *in vivo* loss-of-function studies, such as gene knockdown and gene knockout, are commonly employed to identify and annotate genes. To knock down a gene, RNA interference (RNAi) (1,2) has been mostly adopted where the homology-dependent degradation of the target message RNA (mRNA) occurs due to base-pairing between mRNA and microRNA (miRNA) or small interfering RNAs (siRNA) (3). Correspondingly, a miRNA contains ∼21-24 nucleotide RNAs and is loaded into Argonaute (AGO) protein (4) to form an effector complex known as RNA-induced silencing complex (RISC) (5,6); whereas a siRNA (Figure 1) is a small duplex RNA molecule with ∼21-nt ribonucleotides. The siRNA can be produced *in vitro* by chemical synthesis, or generated *in vivo* via drosha/dicer mediated cleavage of longer duplex RNA precursors (7). Owing to the rapid development of drug delivery materials (8), the RNAi-based therapeutics (9) have experienced remarkable successes in the last couple of years. For example, the first RNAi-based medication Patisiran (Onpattro is its trade name) has been proven by FDA in August 2018 for treatment of polyneuropathy in people with hereditary transthyretin-mediated amyloidosis.

**Figure 1.**
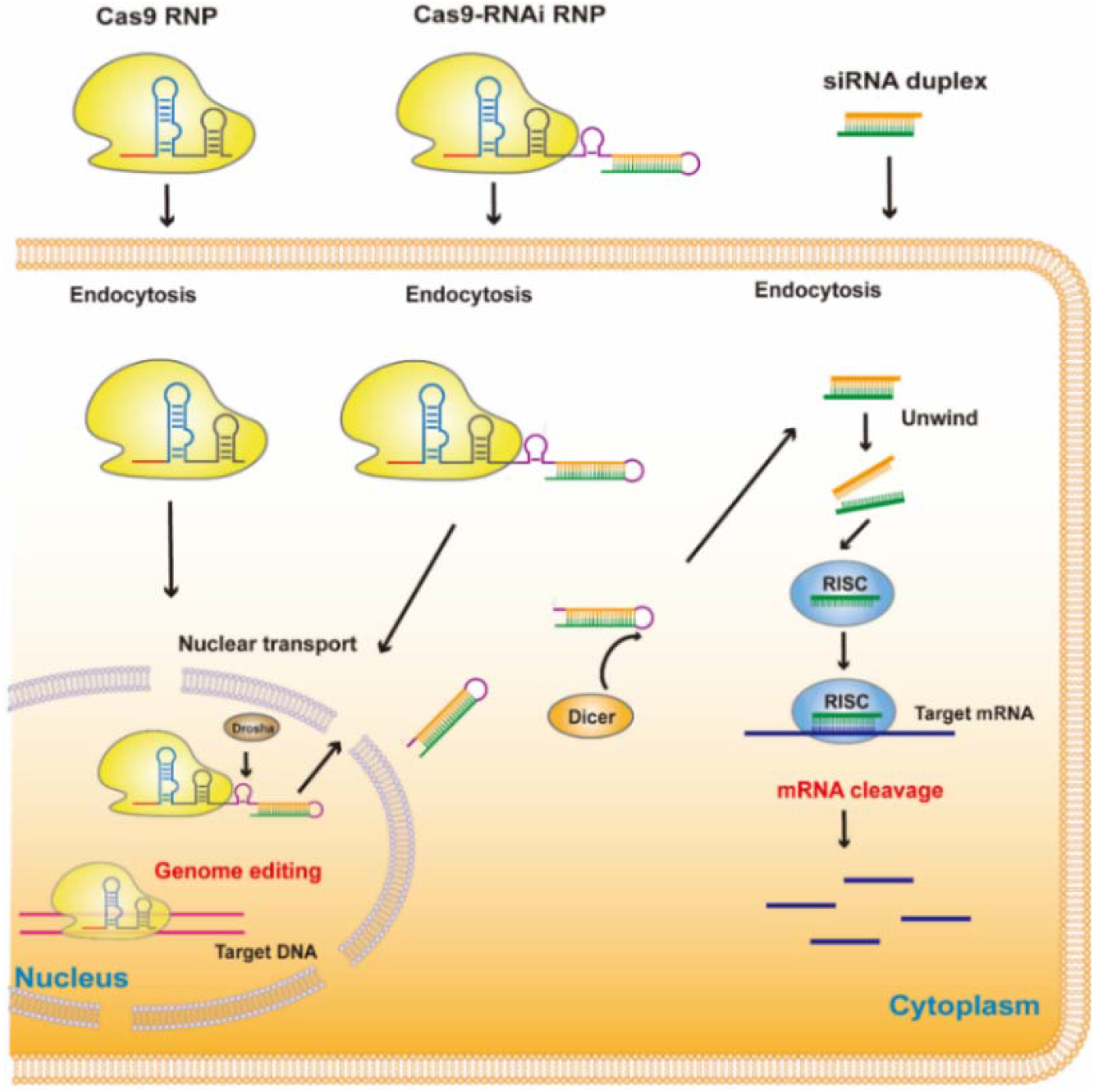
Schematic depiction of transferring Cas9 RNP, Cas9-RNAi RNP and siRNA into a cell for knocking out and/or knocking down the target genes.

To knock out a gene, several genome-editing tools can be chosen, including zinc-finger nucleases (ZFNs) (10), transcription activator-like effector nucleases (TALENs) (11), and clustered regularly interspaced short palindromic repeats (CRISPR) associated nucleases (12,13). Recently, the CRISPR technology has attracted the most attention due to its simplicity, high efficiency, and ease of use. Among the discovered CRISPR/Cas nucleases, Cas9 is most frequently used. This enzyme cleaves the target dsDNA with a gene-specific 20-nt-long complementary region in the single guide RNA (gRNA) (13), producing specific gene deletions via the nonhomologous end joining (NHEJ) pathway, or precise gene repair via the (homologous dependent repairing) HDR pathway. Although the CRISPR toolkits are powerful, limitations still exist such as off-target effects, false positive hits, and low efficiency. Therefore, people have been dedicated to developing new gene-editing tools.

To deliver CRISPR cargos into cells and animals, the virus vectors (e. g. adeno-associated viruses, AAV) are extensively employed. However, researchers have increasingly found that direct delivery of CRISPR/Cas ribonucleoproteins (RNPs) (14-16) for genome editing is a better solution, owing to the advantages (17-19) including reduced off-target effects, low toxicity, and high editing efficiency. Recently, we developed a method (20) for direct preparation of Cas9 RNPs from *Escherichia coli* by co-expression of Cas9 and the target specific single-guided RNAs, achieving convenient and cost-effective production of RNPs. In addition, several reports (21-23) have indicated that extending the 3’ end of gRNA does not impair the activity of Cas9 RNP. For example, Wang et al (21) established a CRISPR-Cas9 platform that can sense miRNA by introduction of a designed gRNA precursor flanked by two extended miRNA-complementary binding sites. Inspired by these works, we aim to develop a gene-editing method that could combine the merits of RNAi and CRISPR technology. In this regard, we designed a gRNA (termed “gRNA-shRNA”) which has a 3’ extension containing a short hairpin RNA (shRNA) (Figure 1). Once being delivered into cells, the shRNA will be processed to a functional siRNA via dorsha/dicer enzyme mediated cleavage (5). We termed this RNP as Cas9-RNAi, and prepared it by our *in vivo* self-assembling method (20). Accordingly, the Cas9 enzymes and the sequence-specific “gRNA-shRNA” ribonucleotides were co-expressed in *Escherichia coli* strain HT115(DE)3. Then, the Cas9-RNAi RNPs were purified and transferred into mammalian cells, achieving knocking out and knocking down the target genes simultaneously. More importantly, we constructed a Cas9-RNAi RNP containing a shRNA specifically against the human *LIG4* gene (24,25), resulting in over 2-fold enhanced HDR-mediated genome editing efficacy. Taken together, we develop a powerful and simple-to-use CRISPR RNP toolkit, enabling multidimensional genome manipulation and showing great potentials for precise genome editing, gene function analysis, and gene therapy.

## MATERIALS AND METHODS

### Construction of the co-expression plasmids of Cas9 RNPs

The co-expression plasmids for all Cas9 RNP variants, including wild type (wt) Cas9, Cas9-RNAi A and Cas9-RNAi B, were constructed according to the protocol described previously (20). To construct the “gRNA-shRNA” (Figure 2), the original gRNA was substituted by the corresponding 3’-extending gRNA. We deposited the plasmid of Cas9-RNAi RNP (Supplementary Figure S1) along with map and sequences in Addgene.

**Figure 2.**
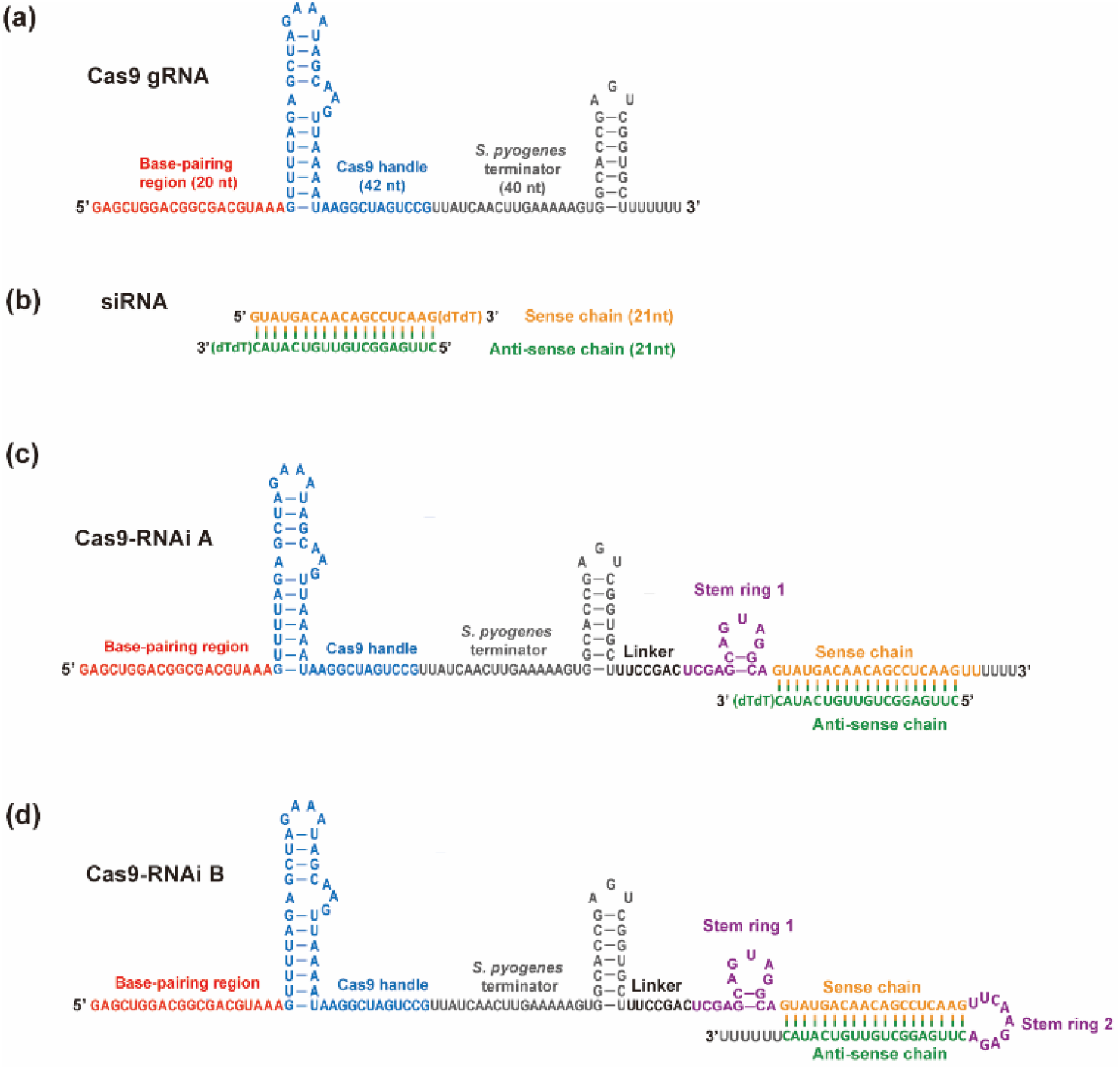
Design of gRNA, siRNA and the “gRNA-shRNA” ribonucleotides. (a) A wt Cas9 gRNA is composited of three regions, including a 20-nt base-pairing region targeting the *GFP* gene, a 42-nt Cas9 handle structure for Cas9 protein binding, and a 40-nt transcription terminator from *S. pyogenes*. (b) A chemically synthesized siRNA duplex against human *GAPDH*. (c) The gRNA of Cas9-RNAi A has a 3’ extension containing a linker, a drosha enzyme cutting site, and a sense chain of siRNA. (d) The gRNA of Cas9-RNAi B has a 3’ extension containing a linker, a drosha enzyme cutting site and a complete shRNA.

### Purification of the Cas9 RNPs by Ni-NTA affinity column

The co-expression plasmids of Cas9 RNPs were transferred into *E. coli* HT115 cells and then cultured at 37°C by using LB medium till OD_600_ reached 0.8. After that, we added 0.5 mM IPTG and turned down the temperature to 16 °C for expression of the Cas RNPs. The *E*.*coli* cells were harvested and lysed in lysis buffer (20 mM Tris-HCl, pH 7.4, 100 mM NaCl). The Cas9 RNPs were loaded onto the Ni-NTA column. Following the standard Ni-NTA affinity purification procedure, the Cas RNPs were eluted with washing buffer (20 mM Tris-HCl, pH 7.4, 500 mM imidazole, 300 mM NaCl). Finally, the Cas RNPs were dialyzed, concentrated, and stored at -80°C in the storage buffer (20 mM Tris-HCl, pH 7.4, 300 mM NaCl, 20% glycerin).

### *In vitro* endonuclease activity assay

The purified Cas RNPs were directly applied to digest the plasmids containing target dsDNA sequences. The digestion reaction was typically carried out in a 10 μL volume of reaction mixture, composed of 1 μL 10× buffer 3.1 (NEB), 200 ng Cas9 RNP, as well as 300 ng plasmids, at 37 °C for 30 min followed by termination of the reaction at 85 °C for 5 min. The cleaved DNA fragmentation was evaluated by 2% agarose gel electrophoresis.

### PAGE analysis of the guide RNA onto RNP

Firstly, the 800 nM of prepared wt Cas9 RNP or Cas9-RNAi RNP was treated with protease K for 10 minutes. Next, the reaction was stopped by mixing the samples with 2xRNA loading dye (95% formamide, 18 mM EDTA, 0.025% SDS and 0.025% bromophenol blue) and heating it for 5 min at 95°C. The products were resolved by 20% denaturing polyacrylamide gel electrophoresis (PAGE), stained with SYBR Gold (Invitrogen), and analyzed by the ImageJ and Origin software.

### Detecting the efficiency of gene silence

The siRNA molecules were ordered from Sangon Biotech (Shanghai, China) and transferred into the corresponding cells by using lipofectamine 3000 (Thermol Fisher, USA). All the Cas9 RNP variants were transferred by using lipofectamine CRISPRMAX (Thermol Fisher, USA). After 48 h, the efficiency of gene silence was determined by RT-qPCR using two-step real-time PCR (Biorad CFX96 real-time system) with Custom Real-time PCR Gene Detection kit (#QPG-010, GenePharma, Shanghai, China) according to the manufacturer’s instructions. Total cellular RNA was prepared using TRIZOL Reagent (Invitrogen, CA, USA). In addition, β-actin was also amplified as an internal control.

We also examined the GAPDH protein expression by western blotting assay. The cells transferred with siRNA or Cas9 RNPs were harvested at 48 h. The total cell lysates and proteins were collected using M-PER mammalian protein extraction reagent (Pierce Biotechnology, Inc., IL, USA), and subjected to sodium dodecyl sulfate-polyacrylamide gel electrophoresis (10%) and then transferred onto poly vinylidene fluoride membrane (EMD Millipore, MA, USA). The membranes were blocked in phosphate-buffered saline containing 5% fat-free milk for 2 h. Then the membranes were incubated with monoclonal anti-GAPDH antibodies (#G8795, Sigma-Aldrich; 1:5000 dilution) overnight at 4°C, following incubation with horserad-ish peroxidase-labeled goat anti-mouse IgG secondary antibodies (Jackson ImmunoResearch, PA, USA) (1:2000 dilution) at 25°C for 2 h. As an internal control, β-actin was also assessed using anti-β-actin (#A5441, Sigma-Aldrich). Band intensities were quantified using with Gel Doc™ XR+ (Bio-Rad). Values were normalized to β-actin level, and then expressed relatively to the non-transfected control.

### Detecting the efficiency of genome editing

The HEK293, HeLa, and RAW264.7 cell lines were from Biovector NTCC Inc (Beijing, China). The efficiency of the NHEJ-mediated genome editing was determined as follows. After delivery of Cas9 RNPs into cells for 5 h, the cells were washed and replaced with DMEM (with 10% FBS and 1% antibiotics) and then allowed to grow for another 48 h. Then, cells were harvested to extract genomic DNA using a QuickExtract genomic DNA isolation kit (Omega). Indel assays were performed using T7 endonuclease-I assay. The data was analyzed using ImageJ.

For illustrating efficiency of the *in vivo* HDR-mediated genome editing by Cas RNPs, we firstly employed a BFP-expressing HEK293 reporter cell line which had been constructed before (16,20). To transfer BFP to GFP, the wt Cas9 RNP or other Cas9 RNP variants were delivered together with a 70 nt donor ssDNA (Supplementary Table S1) by lipofectamine CRISPRMAX (26) (Thermol Fisher, USA). The HDR efficiency was calculated from the flow cytometry data using the GFP cell counts/total cell counts. In addition, we employed TIDER (27,28) (data analysis is available at http://tide.nki.nl) to determine the HDR efficiency of the endogenous genes in cells.

## RESULTS

### Rationally designed Cas9 RNP with a “gRNA-shRNA” component

Employing the pre-assembled Cas9 RNPs for genome engineering has several advantages, such as reduced off-target effects and low toxicity. Here, we sought to develop a technology that could combine the merits of RNAi and CRISPR/Cas9 technology. The schematics of our design is illustrated in Figure 2. As an example, the wild type (wt) Cas9 RNP contains a gRNA (Figure 2a) with a 20 nt base-pairing region targeting the GFP gene in a GFP-expressing HEK293 cell line. Taking advantage of the reporter system, the GFP knockout efficiency can be evaluated by the decrease of green fluorescence. Figure 2b shows the sequence of a 21-nt siRNA against *GAPDH*, a human housekeeping gene. To synchronously knock out *GFP* and knock down *GAPDH*, we designed two Cas9 variants: Cas9-RNAi A (Figure 2c) and Cas9-RNAi B (Figure 2d), respectively. The gRNA of Cas9-RNAi A RNP has a 3’ extension containing the sense chain of GAPDH siRNA, a 7-nt linker, and a dorsha enzyme cutting site (stem ring 1). To form a complete siRNA mimic (Figure 2c), ∼5 times as many synthetic anti-sense chains was added *in vitro*. Once being delivered into cells, the extension of gRNA is cleaved by drosha/dicer enzyme to release a functional siRNA. In contrast, the gRNA of Cas9-RNAi B RNP has a 3’ extension containing a linker, a dorsha enzyme cutting site (stem ring 1), as well as an intact shRNA with a stem ring 2. By this design, the introduced shRNA will be eventually processed to a functional siRNA via dorsha/dicer enzyme mediated cleavage within cells.

### Streamline production of Cas9 RNPs from *E. coli* HT115(DE)3 cells

Recently, we reported a method that enables streamline preparation of Cas9 RNPs in *E. coli* strain BL21(DE3) (20). Compared to the *in vitro* assembling Cas9 RNPs by conventional method (29), the Cas9 RNPs made by this *in vivo* self-assembling approach exhibit significantly enhanced stability and activity. In this work, we harnessed the same strategy to prepare all Cas9 RNP variants (Supplementary Figure S2a), including wt Cas9, Cas9-RNAi A and Cas9-RNAi B. When purified, their nuclease activities against the target plasmids were detected *in vitro* (Supplementary Figure S2b). For a 10 μL reaction system, if ∼300 ng dsDNA can be fully cleaved in less than 30 min at 37°C by 200 ng Cas9 RNPs, the batch of Cas9 RNPs is believed to be qualified for the following cell experiments.

As known, *E. coli* HT115(DE)3 is a RNase III deficient strain which has been applied to the RNAi feeding (30,31) experiment for *Caenorhabditis elegans*. Thus, to prevent the degration of the “gRNA-shRNA” within *E. coli* cells, we selected HT115(DE)3 instead of BL21(DE3) as the expression strain. The result (Supplementary Figure S3) indicates that the complete “gRNA-shRNA” was formed according to our design.

### Multidimensional genome manipulation

To verify the concept in Figure 1, we transferred all three kinds of Cas9 RNPs to a GFP-expressing HEK293 cell line by Lipofectamine CRISPRMAX. The FACS analysis of the decrease of green fluorescence indicates the same-level gene knockout (NHEJ) efficacy of ∼70% for all RNPs (Figure 3a), revealing that extending the 3’ end of gRNA does not affect the activity of Cas9 RNPs. To identify the efficiency of gene silence, we transfected siRNA, Cas9-RNAi A RNP, and Cas9-RNAi B RNP individually to cells. The mRNA level of GAPDH was determined by qPCR at 48 h after transfection (Figure 3b). The similar silence effects were observed for Cas9-RNAi B RNP and siRNA, whereas a less inhibition effect was observed for Cas9-RNAi A RNP.

**Figure 3.**
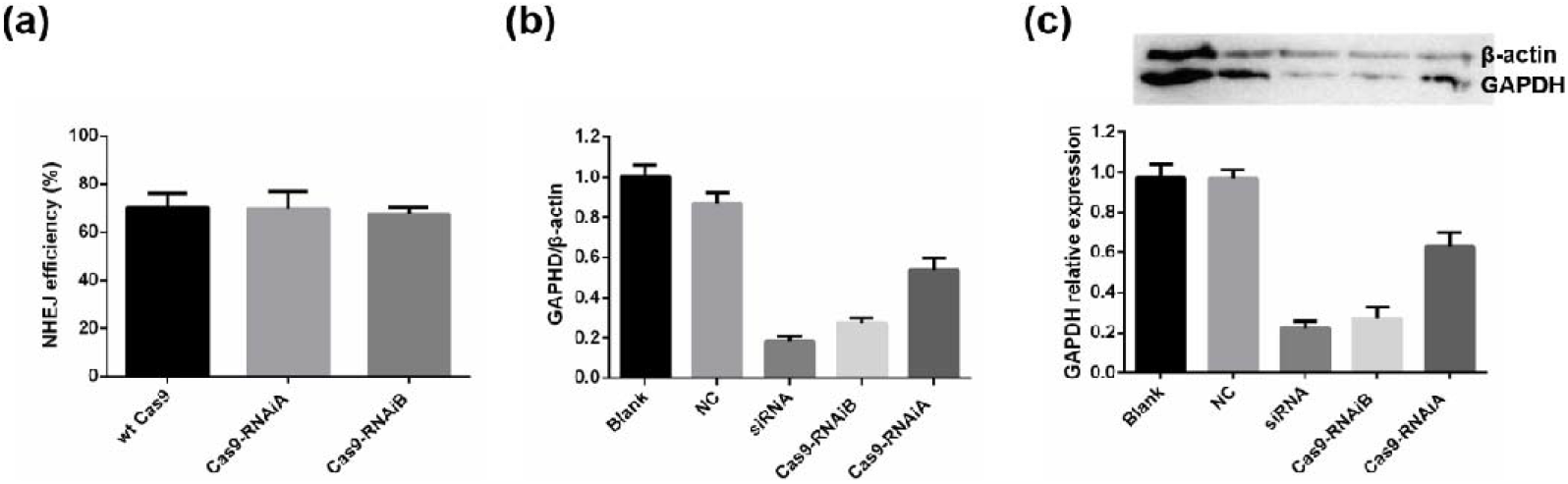
Detecting the efficiencies of gene knockout and gene knockdown. (a) The gene knockout (NHEJ-mediated genome editing) efficiency was measured by FACS analysis at 48 h after transfection; (b) the qPCR analysis of GAPDH at mRNA relative to β-actin at 48 h after transfection; (c) the protein-expression level of GAPDH relative to β-actin at 48 h after transfection. Blank represents the cells without transfection; NC represents the cells that were transfected with scrambled siRNA. Note: the siRNA was transferred by Lipofectamine 3000; all the Cas9 RNPs were transferred by Lipofectamine CRISPRMAX.

Moreover, we measured the efficiency of gene knockdown on the protein-expression level of GAPDH at 48 h after transfection (Figure 3c). Consistent with the qPCR results, the western blot results indicate that both siRNA and Cas9-RNAi B RNP exhibit much higher gene silencing efficacy than Cas9-RNAi A RNP. Perhaps, the synthetic anti-sense chains of siRNA cannot sufficiently bind to the gRNA in Cas9-RNAi A RNP, which leads to obtaining a small number of siRNA mimics. Besides, the anti-sense chains of siRNA should be added to prepare Cas9-RNAi A RNP, which is much more time-consuming and costly. Therefore, Cas9-RNAi B RNP is a much better design. For clarity, we termed it as Cas9-RNAi RNP in the following studies unless otherwise stated.

### Simultaneously knock out and knock down the endogenous genes

The above proof-of-concept experiment has proven that the designed Cas9-RNAi RNP enables multidimensional genome manipulation at DNA and mRNA level in a reporter cell line. To verify the fidelity of the method, we further carried out experiments on different mammalian cells, including HEK293T, and HeLa and RAW264.7.

Firstly, we designed a Cas9-RNAi RNP to knock out human *PRDX4*. This gene encodes peroxidase-4, which plays a regulatory role in the activation of NF-kappaB. In the meanwhile, the *GAPDH* was knocked down. In the next, we prepared the Cas9-RNAi RNPs and delivered them to the corresponding cells. The NHEJ-mediated genome editing capability was evaluated by the T7 endonuclease-I assay (Figure 4), revealing efficient indel gene editing. Meanwhile, the gene silencing effect was determined by RT-qPCR. The results were summarized in Table 1, which indicates that the Cas9-RNAi RNP can simultaneously knock out and knock down the endogenous genes in mammalian cells.

**Table 1.**
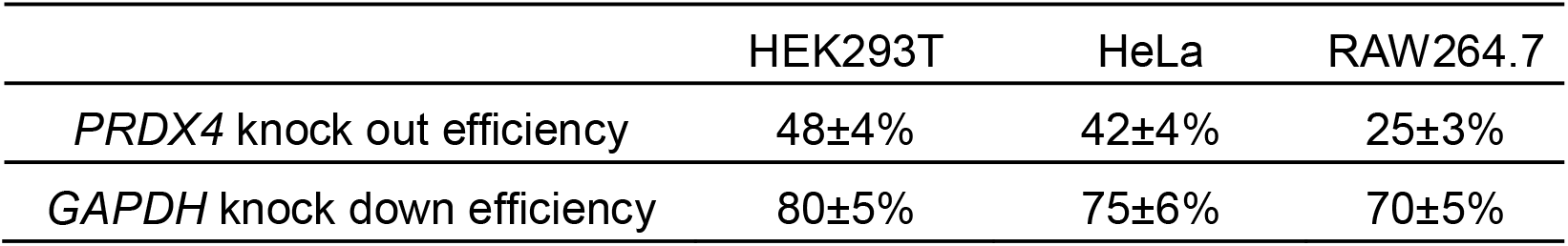
The efficiencies of knocking out and knocking down the genes.

**Figure 4.**
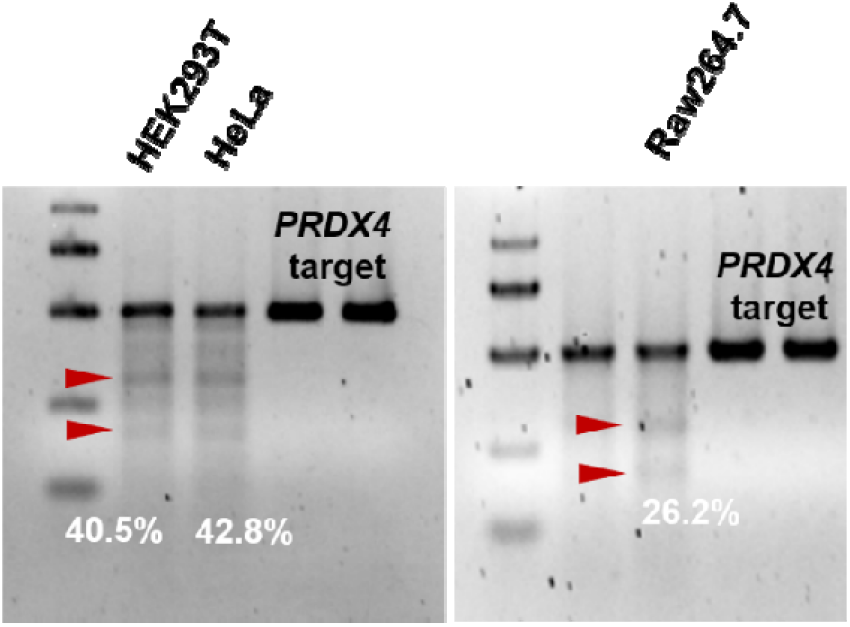
Delivery of Cas9-RNAi RNPs to the target *PRDX4* genes in HEK293T, HeLa and RAW264.7 cells resulted in efficient indel gene editing, as determined by the T7 endonuclease-I assay. The data was analyzed using ImageJ.

### Design of gRNA-shRNA for enhancing the HDR-mediated genome editing

Beyond disrupting genes, the CRISPR/Cas9 tools can be adopted to introduce the gene of interest. In the presence of donor DNAs, precise gene repairing can be achieved via the HDR pathway (14). Previously, several works have demonstrated that inhibition of the key factors in NHEJ pathway, such as Ku70, Ku80 (32) and DNA ligase IV (LIG4) (25) could enhance the efficiency of HDR. Learning from these works, we constructed a Cas9-RNAi (termed Cas9-LIG4) RNP containing a specifically introduced shRNA against *LIG4*. By contrast, we generated a Cas9-NC RNP containing an irrelevant shRNA. Moreover, the wt Cas9 RNPs were delivered into cells together with the synthetic siRNAs (siRNA-LIG4 or siRNA-NC) as the controls.

To measure the efficiency of HDR-mediated genome editing, we employed an engineered blue fluorescent protein (BFP)-expressing HEK293 cell line as the reporter system (16,20). When correct genome repairing occurred after co-delivery of Cas9 RNPs and a 70 nt ssDNA donor (Figure 5a), the BFP-HEK293 cells would be converted to green fluorescent protein (GFP)-expressing cells (Figure 5b). The gene silencing effect of *LIG4* were estimated by RT-qPCR, showing ∼75% inhibition efficiency. The HDR efficiencies were estimated from FACS data (Supplementary Figure S4). The HDR efficiency of Cas9-LIG4 RNP is ∼24±4%, which is ∼2-3 times higher than wt Cas9 RNP (10±2%) or Cas9-NC RNP (9±1%). In addition, the HDR efficiency of wt Cas9 RNP transferred with siRNA-LIG4 is 17±4%, which is much higher than that of wt Cas9 RNP with siRNA-NC (8±1%). Remarkably, transferring Cas9-LIG4 RNP only has the highest HDR efficiency (Figure 5c). In addition, we also adopted the TIDER analysis method (27,28) to detect the HDR efficiency. The similar results were obtained (Supplementary Figure S5), which further confirmed that use of Cas9-LIG4 RNP can significantly improve the homology-directed genome editing.

**Figure 5.**
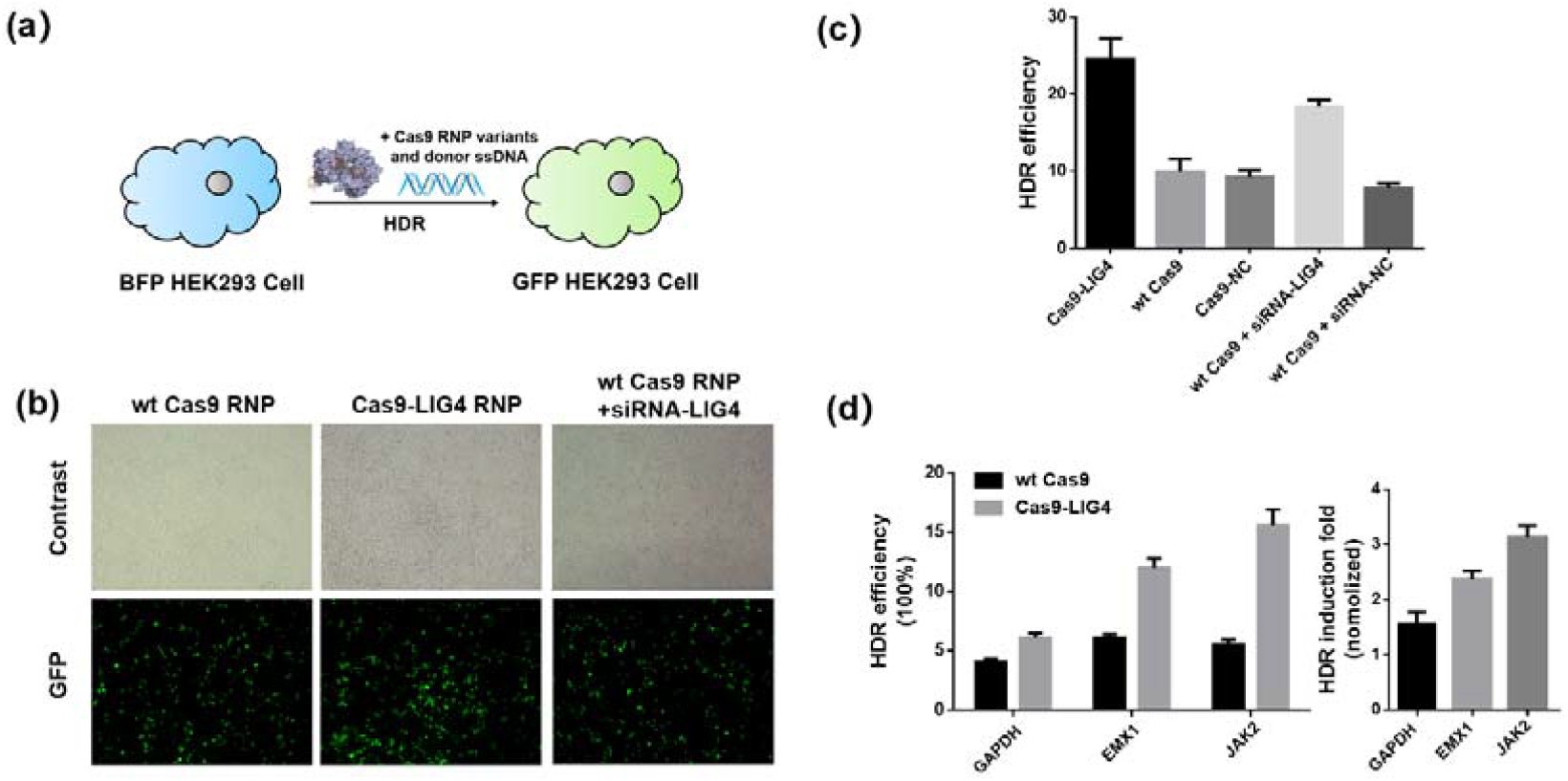
Detection of the HDR-mediated genome editing. (a) Delivery of Cas9 RNPs and donor ssDNA in BFP-HEK293 cells can convert them into GFP-HEK293 cells. (b) The bright-filed and fluorescent images of BFP-HEK293 cells after delivery of wt Cas9 RNPs (left), Cas9-LIG4 RNPs (middle), and wt Cas9 RNPs together with siRNA-LIG4 (right) by lipofectamine CRISPRMAX in 48 h later. All the experiments are performed in the presence of donor ssDNA. (c) The HDR efficiency was determined by GFP expression due to BFP editing according to the flow cytometry data. (d) The HDR efficiency of the corresponding endogenous gene was determined by TIDER analysis method, endonuclease-I assay. Note: data are shown as the mean ± SD.

In the end, we constructed different Cas9-RNAi RNPs to edit the endogenous genes in HEK293T cells, including *GAPDH, EMX1* and *JAK2*. A ssDNA donor (Supplementary Table S1) containing a *Hind* □ cleaving site (Supplementary Figure S6) was applied to measure the HDR efficiency. Also, the TIDER approach was employed to determine the HDR efficiency. As a result, an average ∼ 2-fold HDR efficiency (Figure 5d) was achieved for each gene. Together, we develop a powerful CRISPR/Cas9 RNP tool for precise genome editing via homology-directed repair.

## DISCUSSION

Direct injection of Cas9 RNPs into tissues or organs has been proven as a promising strategy for genetic therapy (17,18,33). However, the most challenge is that the Cas9 RNPs usually have poor stability and limited lifetime within organisms. Therefore, very high dose of Cas9 RNPs must be delivered into patients. Recently, we report a method (20) to timely produce Cas9 or Cas12a RNPs from *E. coli* by co-expression of the Cas enzymes and the target specific single-guided RNAs. Importantly, the prepared Cas RNPs showed significantly enhanced stability, as well as profound HDR-mediated genome editing in mammalian cells. Based on these achievements, here we sought to develop a novel CRISPR/Cas RNP tool.

The gene manipulation technology, such as gene knockdown and gene knockout, is the cornerstone of modern life sciences. Among gene knockdown tools, RNAi (2) is most widely utilized for understanding the function of genes and curing inherited diseases. Regarding gene knockout technologies, the CRISPR/Cas9-associated methods (12) have attracted the most attention in recent years. However, there is no tool to unify these two technologies to date. Thus, here we aim to develop a method that could combine the merits of RNAi and CRISPR.

We refined the designs, and finally constructed a Cas9-RNAi RNP with a “gRNA-shRNA” competent. Compared to original wt Cas9 RNP, this RNP has a 3’-extending functional shRNA that can be converted to siRNA and conduct the function of gene silencing once getting into cells. Besides, it remains the profound genome-editing activity in melamine cells. In other words, we develop a CRISPR/Cas9 RNP tool which enables multiple genome manipulation, such as simultaneously knock down and knock out genes.

In the end, we designed a Cas9-RNAi RNP (named Cas9-LIG4) which has a shRNA specifically against the gene of human *LIG4*. Transferring Cas9-LIH4 RNPs into mammalian cells resulted in significantly improved (over 2-fold) HDR efficiency. In future, the scale prepared Cas9-LIG4 RNPs might be used for genetic therapy. For example, we are developing a method for precisely editing the human CAR-T cells (34) by using the custom-built Cas9-LIG4 RNPs.

In conclusion, we develop a powerful and simple-to-use CRISPR/Cas9 RNP tool, which unifies RNAi and CRISPR and achieves multidimensional genome manipulation. We anticipate that the present method will be applied for precise genome editing, gene function analysis and gene therapy in future.

## DATA AVAILABILITY

The data underlying this article will be shared on reasonable request to the corresponding author.

## SUPPLEMENTARY DATA

Supplementary Data are available at NAR Online.

## ACKNOWLEDGEMENTS

*Author contributions*: W.L.S.: Performed all experiments and data analysis, devised the experimental strategy. W.H.Y.: Assisted with the experiments and data interpretation. L.X.M.: Gave informative suggestions. J.Q.: Contributed to the experimental strategy, assisted with the experimental design and data interpretation. Y.L.: Devised experimental strategy and design, assisted with the data interpretation, and wrote the manuscript.

